# Curbing zoonotic disease spread in multi-host-species systems will require integrating novel data streams and analytical approaches: evidence from a scoping review of bovine tuberculosis

**DOI:** 10.1101/2023.05.08.539893

**Authors:** Kimberly Conteddu, Holly M. English, Andrew W. Byrne, Bawan Amin, Laura L. Griffin, Prabhleen Kaur, Virginia Morera-Pujol, Kilian J. Murphy, Michael Salter-Townshend, Adam F. Smith, Simone Ciuti

## Abstract

**Background:** Zoonotic diseases represent a significant societal challenge in terms of their health and economic impacts. One Health approaches to managing zoonotic diseases are becoming more prevalent, but require novel thinking, tools and cross-disciplinary collaboration. Bovine tuberculosis (bTB) is one example of a costly One Health challenge with a complex epidemiology involving human, domestic animal, wildlife and environmental factors, which require sophisticated collaborative approaches.

**Objective:** We undertook a scoping review of multi-host bTB epidemiology to identify recent trends in species publication focus, methodologies, scales and One Health approaches. We aimed to identify research gaps where novel research could provide insights to inform control policy, for bTB and other zoonoses.

**Results:** The review included 167 articles. We found different levels of research attention across episystems, with a significant proportion of the literature focusing on the badger-cattle-TB episystem, with far less attention given to the multi-host episystems of southern Africa. We found a limited number of studies focusing on management solutions and their efficacy, with very few studies looking at modelling exit strategies. Surprisingly, only a small number of studies looked at the effect of human disturbances on the spread of bTB involving wildlife hosts. Most of the studies we reviewed focused on the effect of badger vaccination and culling on bTB dynamics with few looking at how roads, human perturbations and habitat change may affect wildlife movement and disease spread. Finally, we observed a lack of studies considering the effect of weather variables on bTB spread, which is particularly relevant when studying zoonoses under climate change scenarios.

**Conclusions:** Significant technological and methodological advances have been applied to bTB episystems, providing explicit insights into its spread and maintenance across populations. We identified a prominent bias towards certain species and locations. Generating more high-quality empirical data on wildlife host distribution and abundance, high-resolution individual behaviours and greater use of mathematical models and simulations are key areas for future research. Integrating data sources across disciplines, and a “virtuous cycle” of well-designed empirical data collection linked with mathematical and simulation modelling could provide additional gains for policy-makers and managers, enabling optimised bTB management with broader insights for other zoonoses.

## Introduction

Emerging infectious diseases represent a significant public health concern as they become more prevalent worldwide [1-3]. It is estimated that about 60% of emerging infectious diseases are zoonotic, 72% of which have been estimated to originate from wildlife [2, 3]. In 2019, thirteen different zoonoses had confirmed cases in humans within the European Union [4]. This has been promoted by human technological advances that have caused an exponential growth in global population size and mobility, leading to increased probability of human-wildlife interactions and, therefore, exposure to zoonotic diseases [5, 6]. Zoonotic disease exposure is particularly worrying when combined with the current human population trend and the expected increase in contacts among humans, livestock and other captive animals, as well as wildlife species [6-8].

A multidisciplinary approach and coordinated collaborations between the public health sector, veterinarians, ecologists and wildlife managers have become key to managing existing and to prevent emerging zoonotic diseases [1]. The importance of interdisciplinary approaches is highlighted by the interlinked nature of human, animal and ecosystem health, which led to the new concept of “One World One Health™” [9, 10]. Due to the complexity of zoonotic disease dynamics, the One Health approach offers the most effective means of studying zoonosis thanks to the collaboration of experts in a variety of research fields. Despite such multidisciplinary efforts, the effect of stressors – i.e., direct and/or indirect disturbances such as hunting, habitat loss, and more broadly habitat and climate change – on animal ecology within human dominated landscapes and the potential emergence of zoonotic disease is still understudied [1]. Overall, there is a lack of empirical evidence on how human perturbations can alter disease transmission and the emergence of zoonotic diseases.

Transmission dynamics of zoonoses involve multiple agents including humans and a diverse range of wild and domestic animals. In order to understand the processes behind their transmission, it is essential to clearly disentangle the role of each agent involved [11]. Due to the complexity of disease transmission and the maintenance of infection within multiple wildlife hosts, the individual components of the transmission chain are often studied separately, which can limit our understanding of the subtle underlying effects explaining disease emergence and transmission. Therefore, a holistic approach is essential to develop a complete picture of the transmission dynamics of zoonotic diseases [11], with research on rabies being an outstanding example of how empirical data can be used to elucidate epidemiological dynamics [12].

Evidence from empirical data can be boosted by mathematical simulations, which are powerful tools for predicting disease transmission trajectories [13]. Simulations of disease transmission through compartmental models (e.g., the Susceptible, Infectious, and/or Removed (SIR) model and its variations) have been used in a variety of disease systems, including the recent COVID-19 pandemic. COVID-19, however, is exceptional in the level of global concern garnered and related significant investment in funding, resulting in the generation of large and accessible empirical datasets [14]. Zoonoses are typically more difficult to model due to the lack of empirical data on disease transmission and associated hosts [15]. Mathematical simulations, therefore, offer opportunities to model zoonoses where empirical data are limited. In addition, such simulations allow us to undertake experiments that are currently logistically unfeasible, too costly, too complex or on “unobservable” phenomena [13, 16].

Studying interactions between and within host species as well as the role played by each host in the transmission chain is a powerful tool that can be used to better understand zoonotic disease dynamics. Wild animal interactions are heterogeneous by nature and vary significantly between different populations as well as individuals. Therefore, it is important to account for this variability to understand the mechanisms behind transmission and subsequently be able to predict and control disease spread [7]. This can be achieved by using network modelling, where heterogeneous contacts between animals can be used to simulate disease transmission [7, 15]. Contact networks can also be used to understand and describe the contact structure among animals using social network analysis (SNA) [7]. For example, SNA can be beneficial for disease management since it is able to identify ‘super-spreaders’ (i.e., highly connected individuals) which can then be targeted for vaccination, allowing for a dramatic reduction in transmission [1, 7]. In addition, new research is looking at integrating SNA with molecular epidemiology (phylodynamics) to better estimate transmission pathways and direction of transmission between individuals [17].

Another important tool in epidemiology is the use of statistical modelling to explain and predict disease risk and distribution over space and time [18, 19], identify disease clusters [5] and model host abundance [20]. Spatial analyses make it possible to closely monitor and manage zoonotic diseases [21]. Additionally, using a diverse range of temporal scales can be useful in examining long-term disease dynamics, as well as the short-term effects of environmental variables [22]. As ecological processes occur at different scales (from single study sites to macroecological scales), the spatial scale used for disease distribution modelling is crucial in understanding how these processes exacerbate the spread of zoonotic diseases [23, 24]. Large spatial scales (i.e., global, continental) can examine the broader picture and disentangle how host abundance and abiotic factors influence disease prevalence [11]. Smaller spatial scales (i.e., country, region) can be used to examine population dynamics and pathogen genetic diversity at the local level [11]. Therefore, spatially-explicit analyses can be used to inform One Health approaches, advancing our knowledge when high-frequency data on animal movement are collected. For zoonotic pathogens, consideration of animal movement in the wild (e.g., dispersal and ranging dynamics) or domestic hosts (e.g., intraherd movements across farm habitats or interherd trade movements) at an individual level can provide additional insights into spatial processes. Temporal patterns are important to consider as many zoonotic diseases show seasonal variations (e.g., Zoonotic enteric diseases such as *Salmonella* spp, *Escherichia coli*, *Giardia* spp) as well as possible daily variations (i.e., due to the circadian rhythm of microbes and pathogens as well as chronobiology of wildlife hosts) in their infection patterns [25-27].

Human alterations to the environment can also shape and modify zoonotic transmission [11, 21, 28, 29]. However, we currently lack a framework to fully understand their impact on host transmission dynamics [11]. Current research has shown different ways in which pathogen transmission can be affected. For example, human-driven changes in the environment can modify interactions between hosts, change host and vector densities, and alter host longevity and movement [11, 29]. A study by Castillo-Neyra *et al*. showed that rabies transmission was spatially linked to water channels, which act as ecological corridors connecting multiple susceptible populations and facilitating pathogen spread and persistence [29]. With cities expanding and providing urban corridors to wildlife, pathogen persistence could become even more of an issue [29], confirming the importance of studying the effect of human perturbations on animal ecology and related implications in disease ecology.

Understanding the epidemiological dynamics of zoonosis is critical from both a health and economic perspective. A critical example is Zoonotic Tuberculosis (zoonotic TB), which was estimated in 2016 to be linked to 147,000 human cases and 12,500 deaths worldwide [30]. Zoonotic TB is driven mainly by *Mycobacterium bovis* (i.e., the causative agent of Bovine Tuberculosis – known also as bovine TB or bTB), which is transmitted by several wildlife hosts and livestock. Britain and Ireland, as well as many other countries worldwide [30], have been increasingly impacted by bTB, resulting in significant economic loss. In Ireland, for instance, 4.2% of cattle herds tested positive for bTB in 2020, leading to the humane killing of 21,467 cattle [31]. This is in addition to the economic costs associated with the national bTB eradication program with €92 million spent in 2018 alone [32]. Similar trends can be observed in the UK, with £70 million spent annually for bTB prevention and control [33]. This disease also raises welfare concerns for wildlife hosts, especially considering its high prevalence in the wild. Badgers *Meles meles*, for example, have been shown to have a bTB prevalence exceeding 40% in hotspot areas in Ireland [34], and red deer in Spain have been estimated to have a prevalence of up to 50% [35].

Bovine TB eradication is prioritised by governments and researchers due to the significant health concerns and economic (trade) impacts. Despite decades of control efforts in several countries, the pathogen has successfully avoided eradication. There are complex reasons as to why this is the case [36], but a primary factor relates to its complex dynamics of transmission and maintenance across differing hosts and the environment. Therefore, new thinking may be required to further investigate if disease control can be driven toward eradication. Detecting gaps in the current bTB literature is an essential step required to identify target areas for future research and to further hone government eradication strategies. Here, we aimed to uncover empirical and methodological gaps in the peer-reviewed literature on bTB. Our intention is to use bTB as an example of a complex multihost zoonotic disease for which recent developments with sampling design, animal monitoring tools and technology, and mathematical modelling has helped to fill the gaps in knowledge and improve our understanding and ability to combat zoonotic diseases more generally.

To achieve our goal, we developed a scoping review of bTB multihost epidemiology focusing on 18 research questions (reported in **Table 1** and conceptually summarised in **Fig. 1**) regarding the type of study, whether, which and how wildlife species have been monitored, what kind of sampling designs and methodological approaches have been used, and whether epidemiological empirical data have been collected. We then gathered data from the peer-reviewed literature on the mechanisms driving inter-and intraspecies bTB transmission, looking in particular at novel and multi-disciplinary approaches, therefore focusing on the scientific literature published in the last six years (January 2017 -July 2022). Our goal is that our work will spark renewed discussion on how to monitor and deal with zoonotic diseases, direct future research, and stimulate focused funding efforts.

**Fig. 1:**
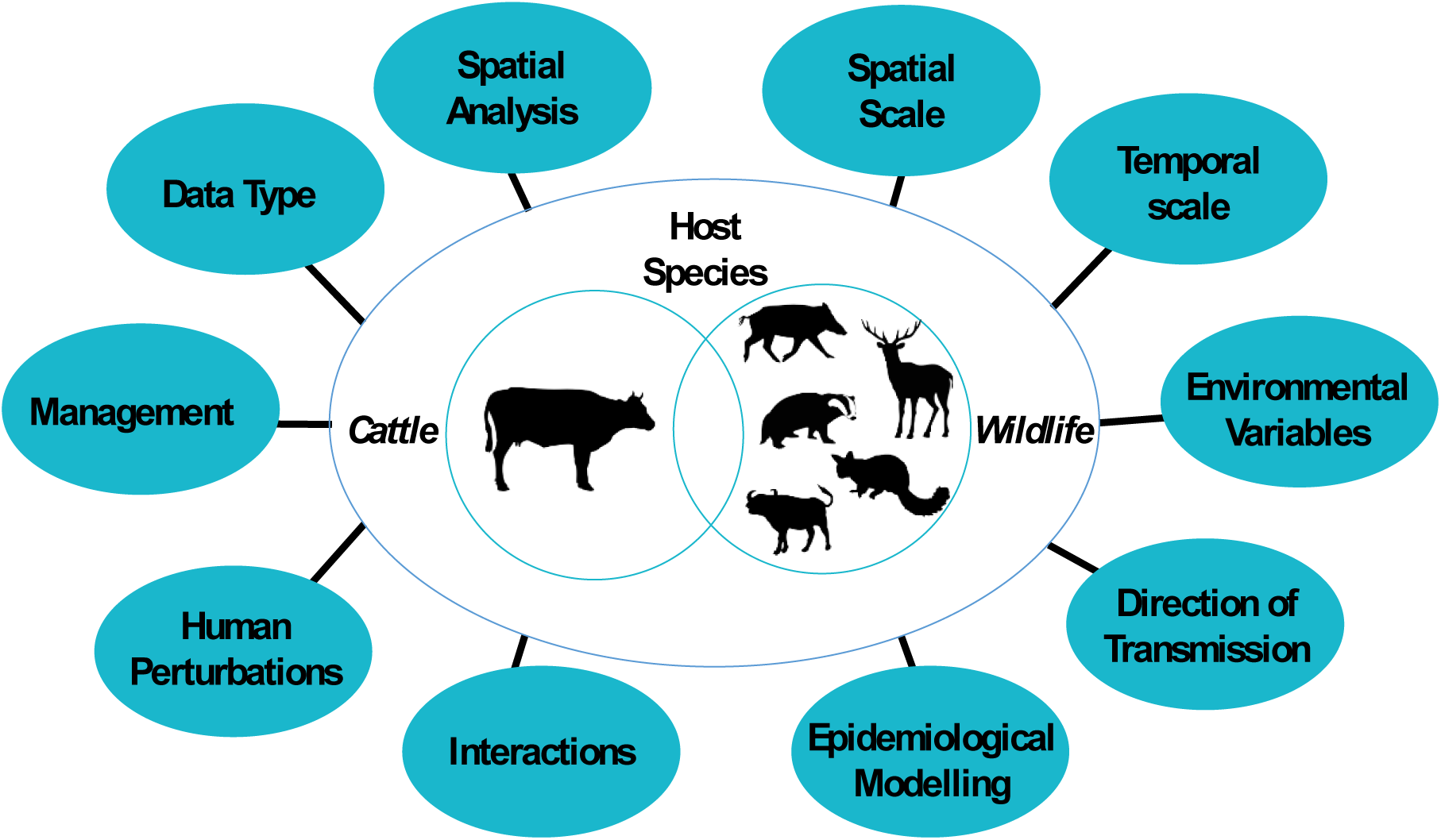
Key host species and topics of interest we screened for in the bovine tuberculosis scientific literature published in the period between 2017-2022. bTB host species include cattle as well as a range of wild species: badger, wild boar, cervid species (with the following species identified in the literature screened: white-tailed deer, red deer, fallow deer, roe deer and wapiti elk), brush-tailed possum and wild buffalo. The circles on the outside illustrate the key information sought in peer-reviewed papers dealing with bTB, which has been expanded and clarified in **Table 1**: type of data collected by researchers; whether spatial analyses were carried out (i.e. in cattle and or wildlife); what type of spatial and temporal scales were considered; whether environmental variables were taken into account (i.e. environment in the farm, environment around the farm and/or climate variables); whether the methodological approach captured the direction of disease transmission; whether the study used common epidemiological modelling techniques (i.e. compartmental models, transmission rates), or whether the study included intra/interspecies interactions in their methodology (i.e. what type of interactions did they look at – e.g. direct and/or indirect, what type of equipment was used to get interactions data and what methodology was used to analyse the data); finally, if human perturbations (i.e. forest felling, culling, vaccination) were taken into account when looking at variables affecting bTB spread, and management solutions to offset the spread of bTB, if any. Animal silhouettes were downloaded from PhyloPic (https://www.phylopic.org/). Cattle, cervid, brushed-tailed possum and wild boar silhouettes are under: CC0 1.0 Universal (CC0 1.0) Public Domain Dedication. Buffalo silhouette is by Jan A. Venter, Herbert H. T. Prins, David A. Balfour & Rob Slotow (vectorized by T. Michael Keesey) under: Attribution 3.0 Unported (CC BY 3.0), link: https://creativecommons.org/licenses/by/3.0/. Badger silhouette is by Anthony Caravaggi under: Attribution-NonCommercial-ShareAlike 3.0 Unported (CC BY-NC-SA 3.0), link: https://creativecommons.org/licenses/by-nc-sa/3.0/.

**Table 1:**
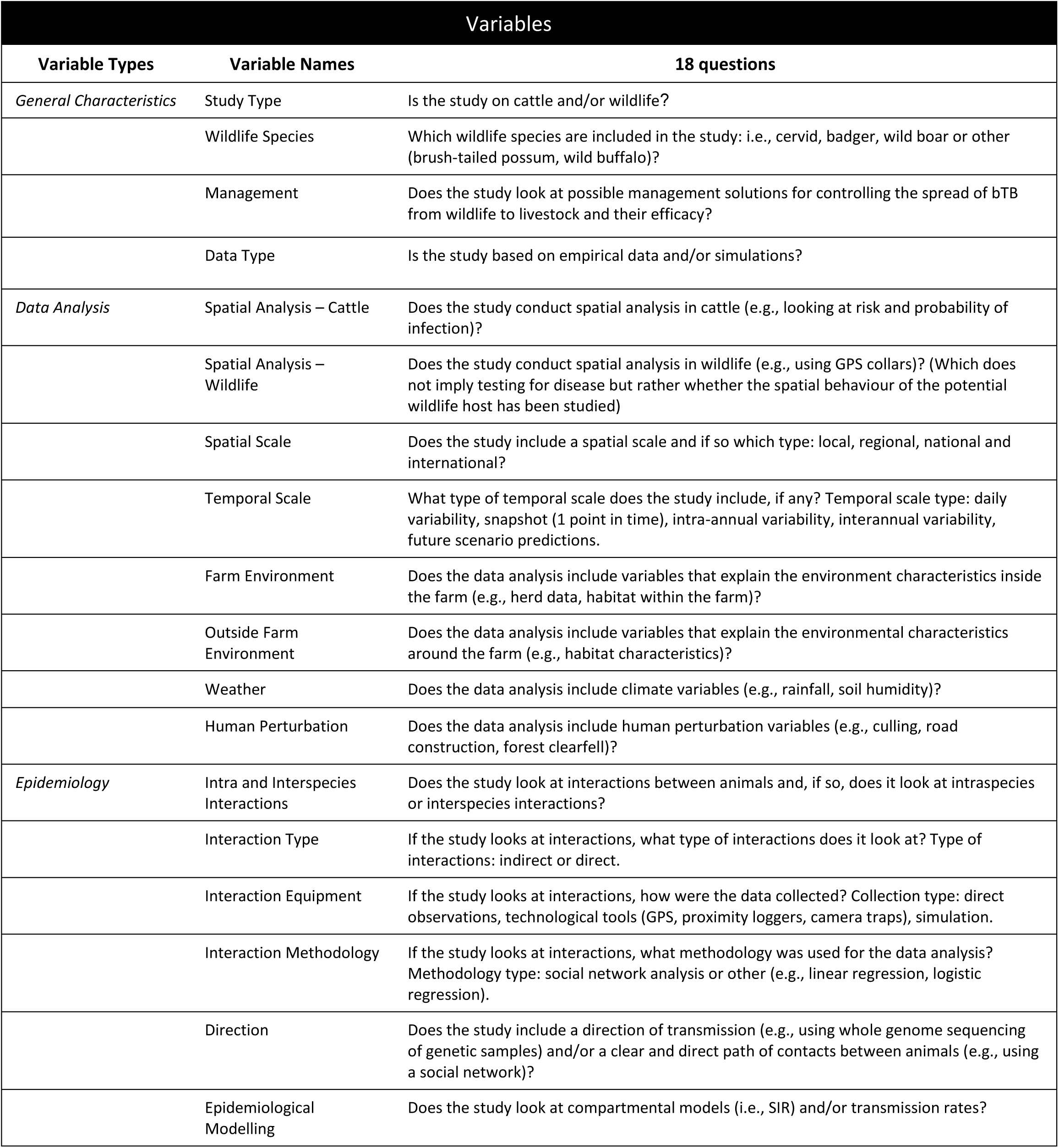
Description of the 18 questions (and related variables) used to screen bTB papers published in the period between 2017-2022. The variables have been divided into three groups depending on the question type. Variables looking at general aspects of the paper (e.g., type of host species included in the study) were grouped as *general characteristics*. Variables concerning the type of data analysis conducted in the studies were grouped as *data analysis*. Finally, variables focused on the type of epidemiological analysis conducted in the papers of interest were grouped as *epidemiology*.

## Review

## 1. Methods

We conducted a scoping review (as per PRISMA guidelines) [37] by sourcing peer-reviewed papers using Web of Science (Clarivate, 2021 Online Version) focusing on bovine tuberculosis, and more specifically its most common cause, *Mycobacterium bovis*, in cattle and several key wildlife hosts. The search terms and list of articles have been summarised in **Additional file 1**. We identified 3,531 potentially relevant papers (i.e., the search included all years of publication) which were uploaded and screened for duplicates using EndNote (Clarivate, Version 20.1.0.15341). Relevant articles were then selected using a PEO (Population, Exposure, Outcomes) eligibility criterium structure [38]. The aim of the PEO is to identify articles of interest by selecting the “Population” (i.e., the subject being affected by the disease/health condition) for a particular “Exposure” (i.e., a disease/health condition) and either a particular “Outcome” or “Themes’’ to examine [38, 39]. The PEO eligibility criterium was chosen since it was in line with the recommendations given for scoping reviews that target literature on etiology and risk factors, such as a particular disease. We decided to use a modified version of the PEO framework structure which also includes themes of interest as potential “Outcomes” [39], as summarised in **Table 2** to guarantee reproducibility. All papers that did not meet the eligibility criteria listed in **Table 2** were removed. bTB has received significant attention in the last few years due to its financial impact on rural economies and the risk it poses to human health. We selected the most recent papers published in the period January 2017 - July 2022 (see **Fig.2** for a summary of the step-by-step process), making up 30% of all relevant papers identified, to capture the most recent methodological approaches used to monitor and combat bTB. The papers were screened to answer the questions of interest summarised in **Table 1**, stored in an excel spreadsheet. The results were then imported and plotted using *<u>ggplot2</u>* in R version 4.1.1 [40].

**Fig. 2:**
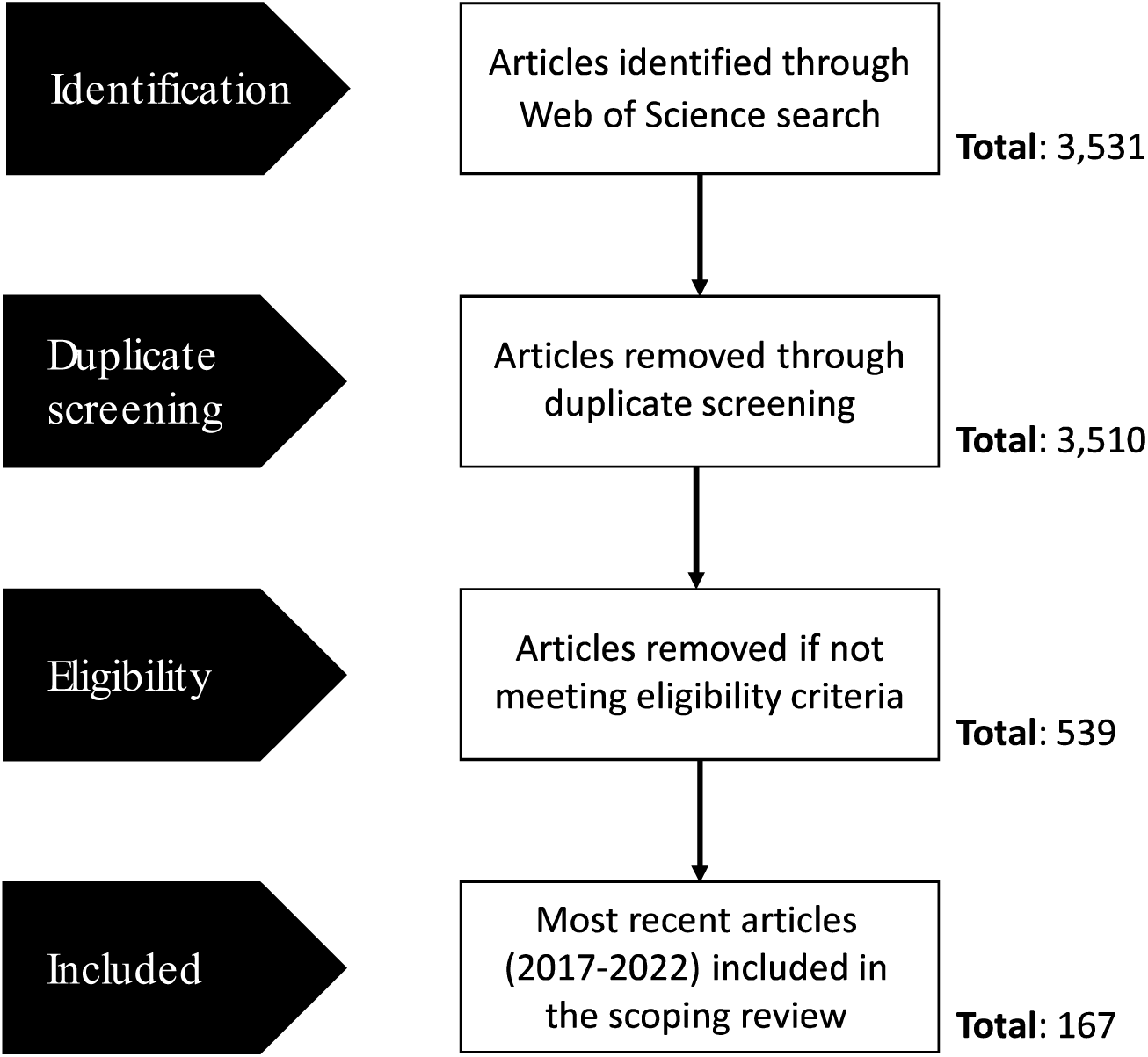
Cascade diagram of the process used in the selection of relevant papers.

**Table 2:** Eligibility criteria used in the selection of papers for the systematic scoping review. The criteria follow a PEO (Population, Exposure, Outcomes/Themes) structure.

| Eligibility Criteria |  |  |  |
| --- | --- | --- | --- |
| PEO Criteria | Explanation | Scoping Review Criteria | Eligible Paper |
| *Population* | Who is the focus of the scoping review? | Cattle, badger, deer, elk, wild boar, wild buffalo, brushed-tailed possum (i.e., known bTB hosts) | Contains at least one of the species of interest as a study species. |
| *Exposure* | What issue is the focal point of the scoping review? | Bovine tuberculosis (i.e., *Mycobacterium bovis*) | Focal point of the paper must be bovine tuberculosis, more specifically the most common cause: *Mycobacterium bovis*. |
| *Outcomes/Themes* | What themes, in relation to the issue, will the scoping review focus on? | ● **Risk:** Does the study assess or evaluate potential risks of spreading bTB and its drivers (biotic and/or abiotic factors)?<br>● **Management:** Does the study look at potential management solutions for controlling the spread of bTB from wildlife to livestock and their efficacy?<br>● **bTB occurrence:** Does the study look at the presence of bTB in cattle and/or reservoir hosts at a spatial and/or temporal level?<br>● **bTB spread:** Does the study look at the mechanisms behind the active spread of bTB between individuals of the same species and/or interspecies transmission? | Focuses on at least one of the themes of interest. |

## 2. Results

Our results are based on 167 peer-reviewed papers published between January 2017 and July 2022. The study location of the papers was representative of 6 continents and 37 different countries (**Fig. 3**). The continent with the highest number of studies on bTB is Europe (n = 103, 52 of which were from the UK), significantly higher than those carried out in much larger continents such as Africa, Asia, and both Americas (**Fig. 3**). We screened all papers for 18 different variables (addressing our 18 questions, see **Table 1**) which we summarised in the following section under the heading: 2.1 **general characteristics** (Sub-headings: “Study species and wildlife species”; “Management and data type”), 2.2 **data analysis** (Sub-headings: “Spatial analysis, spatial scale and temporal scale”; “Farm environment and human perturbations”), and 2.3 **epidemiological analysis** (Sub-headings: “Intra-and interspecies interactions”, “Direction of transmission and compartmental models”). Note that most plots presented below have a sample size of n = 167, corresponding to the number of papers screened, with a few exceptions where this sample size is higher (for example, in relation to species included in the study, because a paper reported data about both wildlife and cattle, therefore contributing to multiple levels of a category) or lower (for example, in relation to epidemiology, where variables of interest were analysed only in the subset of papers describing studies that included epidemiological interactions).

**Fig. 3:**
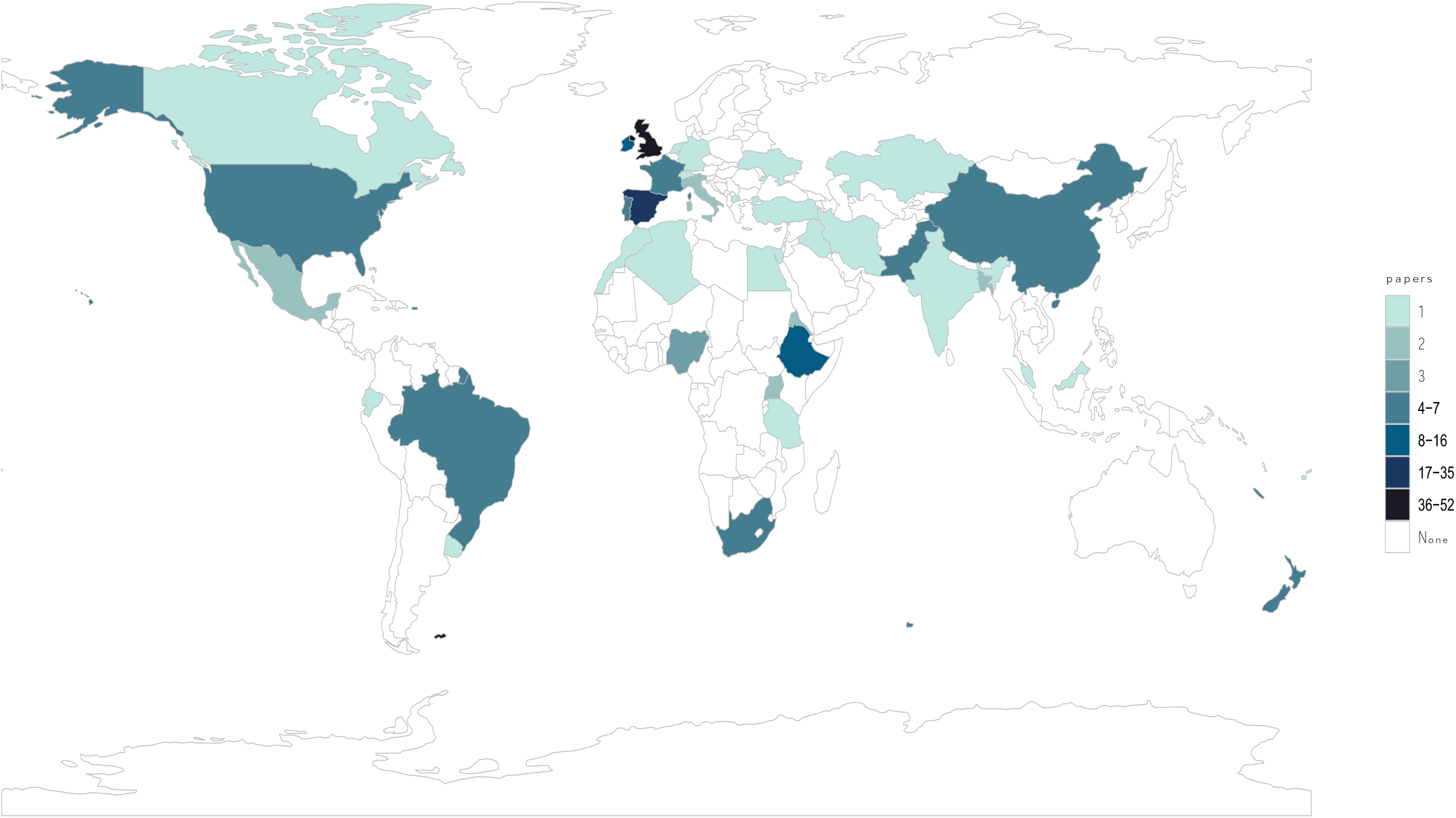
World map showing number of papers screened per country. Number of papers per continent: Europe (103), Africa (26), Asia (19), South America (11), North America (8), Oceania (7).

### 2.1. General Characteristics

#### 2.1.1. Study species and wildlife species

We found that 55% of bTB papers focused on cattle only, whereas 26% of them included both cattle and wildlife species and 19% targeted only wildlife species (**Fig. 4a**). Among those papers reporting wildlife data, we found that the European badger attracted most research effort (60% of wildlife studies), followed by wild boar (29%) and cervid species (24%: 17% red deer, 7% fallow deer, 5% white-tailed deer, 1% roe deer 1%, and 1% wapiti elk, from hereinafter referred to as simply elk) (**Fig. 4b**). Other species were included in 12% of wildlife papers (brush-tailed possum: 8%, wild buffalo: 4%).

**Fig. 4:**
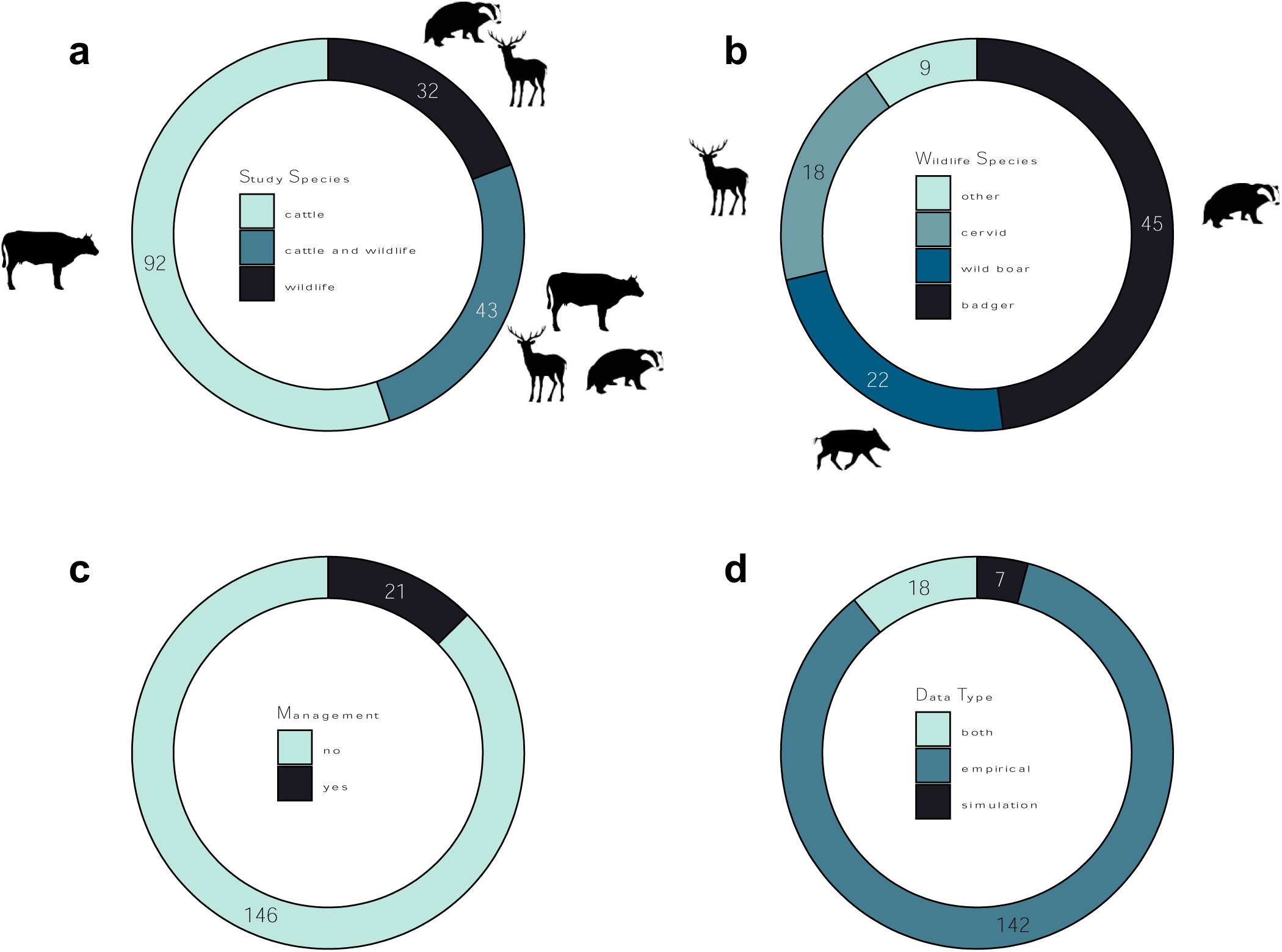
Number of papers screened and reporting data on **a)** study species type (whether the study was on cattle and/or wildlife), **b)** wildlife species, **c)** management (whether a paper investigated potential management solutions and their efficacy), **d)** and data type. Animal silhouettes were downloaded from PhyloPic (https://www.phylopic.org/). Cattle, cervid, brushed-tailed possum and wild boar silhouettes are under: CC0 1.0 Universal (CC0 1.0) Public Domain Dedication. Buffalo silhouette is by Jan A. Venter, Herbert H. T. Prins, David A. Balfour & Rob Slotow (vectorized by T. Michael Keesey) under: Attribution 3.0 Unported (CC BY 3.0), link: https://creativecommons.org/licenses/by/3.0/. Badger silhouette is by Anthony Caravaggi under: Attribution-NonCommercial-ShareAlike 3.0 Unported (CC BY-NC-SA 3.0), link: https://creativecommons.org/licenses/by-nc-sa/3.0/.

#### 2.1.2. Management and data type

Our results highlighted that only 13% of the studies dealt with management solutions (**Fig. 4c**). Management strategies mainly included vaccination (4%) or culling (2%), with 1% looking at both vaccination and culling. We also found that most papers gathered original empirical data (85%), and papers using simulations were very limited (4%), with a remaining 11% of papers combining empirical data and simulations (**Fig. 4d**).

### 2.2. Data Analysis

#### 2.2.1. Spatial analysis, spatial scale and temporal scale

We found that the majority of papers did not include any spatial analysis. Those that did largely focused on spatial patterns in cattle (31%, **Fig. 5a**) slightly more than wildlife (25%, **Fig. 5b**). Among the 51 papers that investigated spatial analysis in cattle, 47% looked at bTB risk and probability of infection; 20% investigated cattle movement outside the farm, 12% looked at cattle interactions with wildlife, 10% analysed the distribution of bTB positive biological samples, and 8% focused on detection rate by spatial unit (e.g., 1 km^2^ grid). Movement inside the farm and interactions of farm animals with both wildlife and cattle were included in 2% of papers. Among the 41 studies which investigated spatial behaviour in wildlife (Fig 5b), analysis was undertaken using a variety of methodologies; direct observations (34%), satellite GPS telemetry (32%), camera traps (7%), genetic samples (12%) and proximity loggers (5%), or spatial patterns predicted by future scenarios modelling or mathematical simulations (12%).

**Fig. 5:**
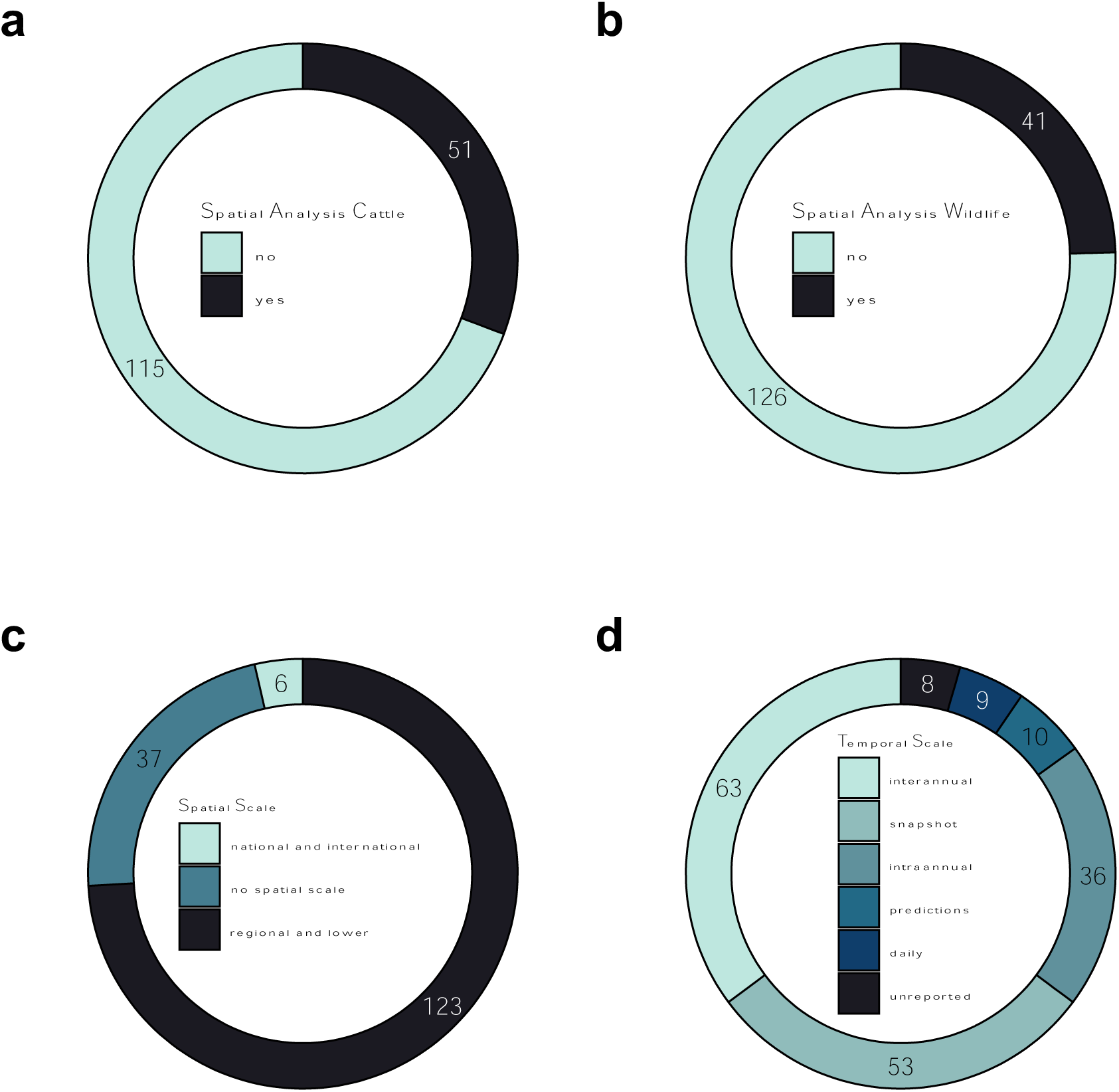
Number of papers screened and reporting data on **a)** spatial analysis of cattle (whether the study included any type of spatial analysis), **b)** spatial analysis of wildlife, **c)** spatial scale, and **d)** temporal scale.

We also found that most papers included spatial scales at the regional level or smaller (74%), with less than 4% papers looking at national and/or international spatial scales (**Fig. 5c**). In regard to temporal scales, 35% of the studies considered interannual variability, whereas 20% tackled intra-annual variability, with a small fraction (9%) of studies including both intra-and interannual variability. Thirty percent of the studies did not analyse any intra-or interannual temporal variability (**Fig. 5d**). Only 5% of the studies looked at fine-scale variability (e.g., days), whereas in a few instances the year of study was not reported at all (5%). Finally, only 6% of papers included predictions for temporal patterns into future scenarios.

#### 2.2.2. Farm environment and human perturbations

When looking at farm characteristics, 44% of the studies included some type of herd data (e.g., herd size, bTB history), with 40% not including any type of in-farm environmental variables (**Fig. 6a**) and 15% of papers incorporating other types of farm characteristics. These included environmental conditions on the farm (e.g., natural habitats, land-fragmentation; 7%), farm location in respect to other farms (6%) and farm location in respect to wildlife (5%).

**Fig. 6:**
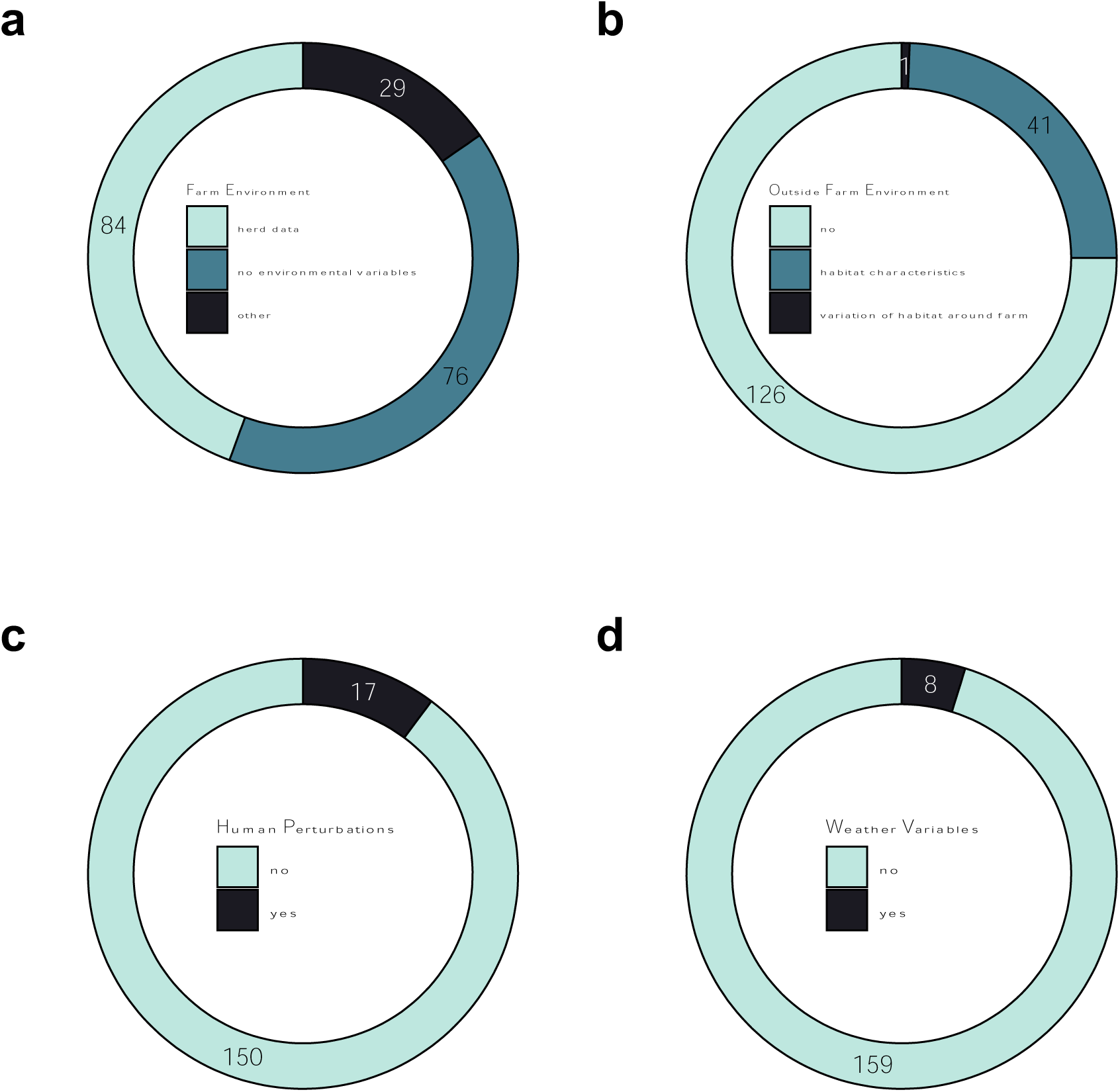
Number of papers screened and reporting data on **a)** in-farm environment (whether the paper analysis included variables explaining environmental characteristics inside the farm), **b)** outside farm environment, **c)** human perturbations (whether the paper analysed the effect of human disturbances on bTB transmission dynamics), and **d)** weather variables.

Environmental conditions outside the farm were included in 24% of the papers’ data analysis (**Fig. 6b**). These studies mainly looked at habitat characteristics around the farm (e.g., wildlife presence, natural habitats), with one paper also including variables focusing on habitat variation (e.g., forest clearfell, new artificial plantations). We also looked at weather variables (e.g., temperature, rainfall) and observed that 5% of papers included these as part of their analysis (**Fig. 6d**). Finally, 10% of the papers screened included human perturbation variables with the vast majority looking at the effect of vaccination and culling on transmission dynamics (**Fig. 6c**).

### 2.3. Epidemiological analysis

#### 2.3.1. Intra-and interspecies interactions

We found that most papers (74%) did not include an analysis on interactions, with 17% of papers looking at intraspecies transmission and 9% at interspecies transmission (**Fig. 7a**). Among the interaction studies, 75% included direct interactions and 25% included indirect interactions (**Fig. 7b**). In addition, interaction data were mostly collected using technological tools (43%; e.g., GPS, proximity loggers, camera traps), followed by direct observations (39%) and a minority using simulations (11%) (**Fig. 7c**). The methodology used to analyse interaction data also varied between papers with most papers using social network analysis (57%), and the others (48%) using a variety of statistical techniques (e.g., linear models, logistic models, differential equations) (**Fig. 7d**).

**Fig. 7:**
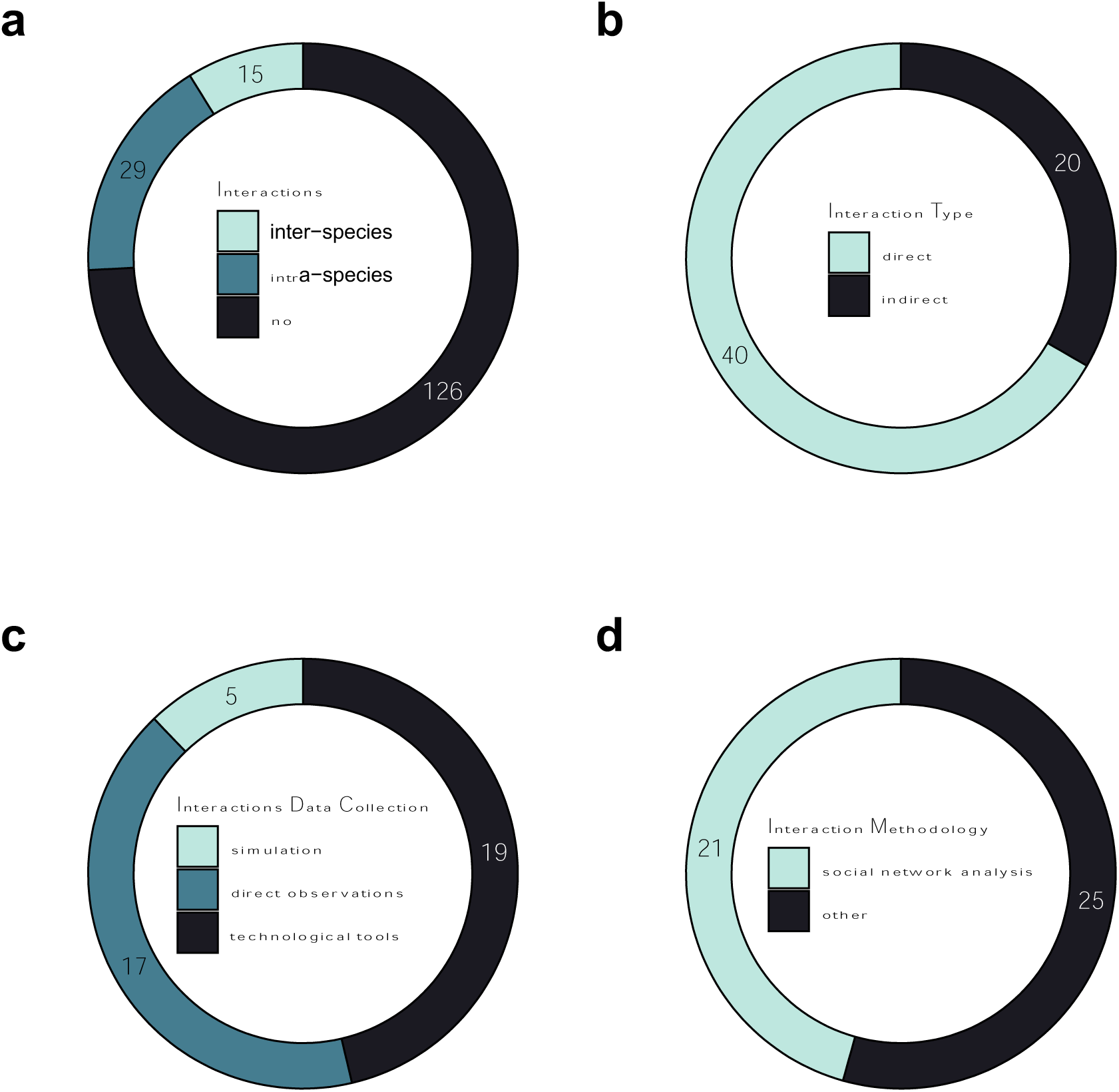
Number of papers screened and reporting data on **a)** interactions (inclusion of interaction analysis i.e., intra-and/or interspecies interactions), **b)** interaction type, **c)** the way interactions were monitored, and **d)** interaction data analysis statistical approach.

#### 2.3.2. Direction of transmission and compartmental models

We found that a limited proportion of the papers (12%) included direction of transmission in their analysis (**Fig. 8a**). We also found that epidemiological modelling techniques (e.g., compartmental models and transmission rates) were adopted in 11% of the studies (**Fig. 8b**).

**Fig. 8:**
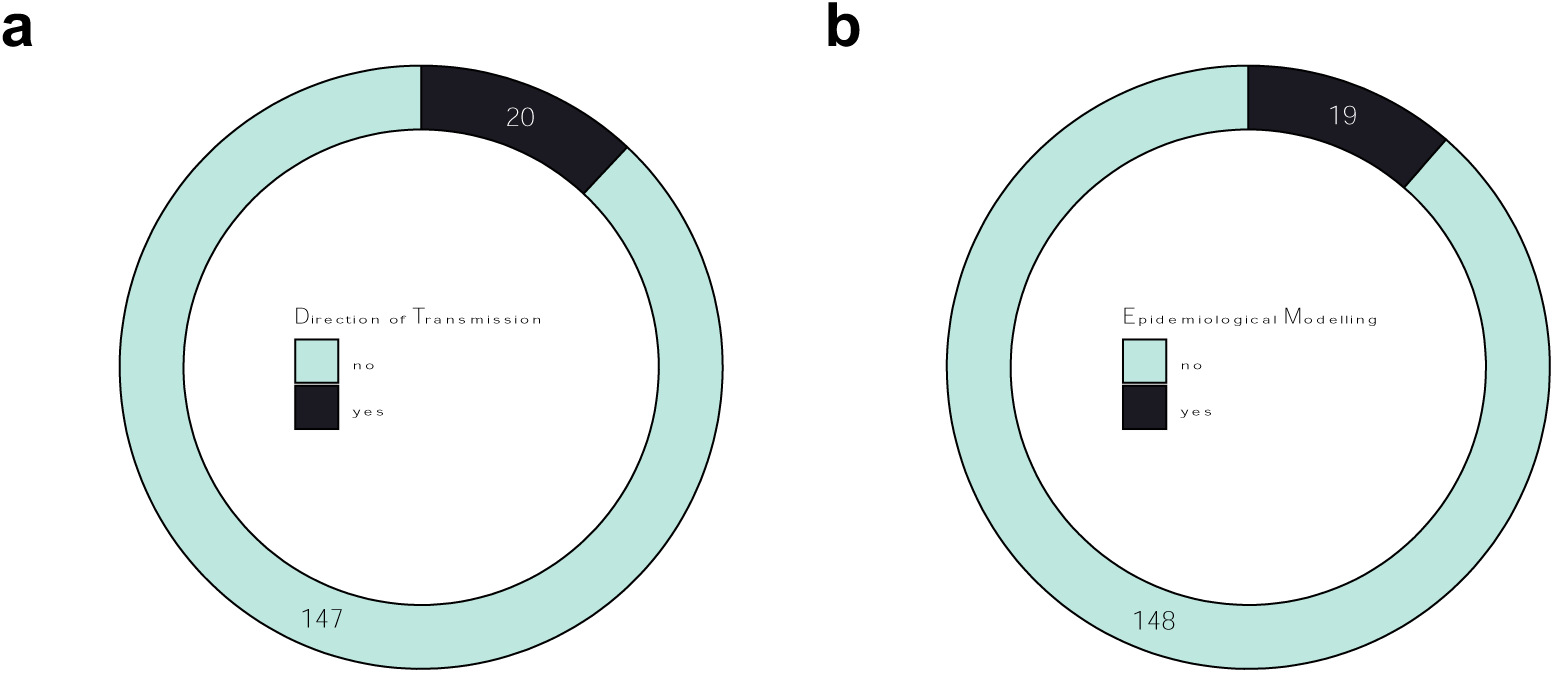
Number of papers screened and reporting data on **a)** direction of transmission (whether this was analysed in the paper, e.g., transmission across species), **b)** epidemiological modelling (i.e., papers included compartmental models and/or transmission rates in the analysis).

## 3. Discussion

In this review we examined bTB papers published between January 2017 to July 2022 corresponding to a third of all bTB papers ever published meeting our selection criteria, highlighting the high level of interest on this zoonosis by researchers aimed at curbing its spread and economic impacts. We found that there has been a great deal of research effort focusing on the badger-cattle bTB episystem; we acknowledge, however, that we found a very limited number of studies on other episystems [41-43]. Our spatially-explicit overview of bTB research efforts (**Fig. 3**) highlights how the badger-cattle episystem has been the focus of research across Europe and particularly in Britain and Ireland. Our review also shows that there has been a different level of attention across episystems -e.g., there has been huge investment of effort and money, resulting in huge output on the badger-cattle-TB episystem, but far less on the multi-host episystems of southern Africa, and we believe we have more to learn from these other systems which are chronically understudied.

Our scoping review found a limited number of studies focusing on management solutions and their efficacy, with very few studies looking at modelling exit strategies [44, 45]. This is not surprising considering the scarcity of studies using mathematical simulations, not only to better understand and predict possible outputs of management solutions, but also to explore long-term bTB dynamics under different scenarios, e.g. [46, 47]. Surprisingly, only a small number of studies have looked at the effect of human disturbances on the spread of bTB involving wildlife host species, and this knowledge gap indeed needs to be tackled in light of research showing the significant role of human perturbations on zoonotic outbreaks and spread [48-50]. Most of the studies we reviewed have focused on the effect of badger vaccination and culling on bTB dynamics with only three studies looking at how roads or human perturbations may affect these dynamics [51-53], and one focusing on the effect of habitat change (clearfell forest operations) on bTB breakdowns [54]. The latter, nevertheless, lacked empirical data on wildlife, which has been indicated as one of the drivers of bTB outbreaks in cattle farms [54]. Finally, we observed that there is a lack of studies looking at the effect of weather variables (i.e., rainfall, soil humidity, temperature etc.) on bTB spread [55]. This is especially important when thinking about wildlife-cattle transmission since it is now thought to occur also through environmental sources [56].

We have carefully evaluated the outcome of our scoping review, and in the following sections we have summarised data types and methodological approaches which, we believe, could contribute to gaining further insights into bTB epidemiology. Based on our review, we have identified a significant gap when it comes to prediction and simulation models, which would be a useful tool for managers to assess disease risk under different land use and climate change scenarios. One major gap that we have to stress is the lack of integration between empirically informed tactical (short-term decision support) and strategic (larger spatial scales and longer term) models blending multiple approaches (though note exceptions before the screening period of our scoping review, see for instance Brooks-Pollock *et al*. [57]). Future research should include compartmental models fitted across space, linked via meta-populations and/or real-landscape multi-host episystems; or agent-based models (ABMs) with empirical data feedback loops. We describe such modelling approaches and their prerequisites in the following sections, beginning with data and monitoring programs, and we continue with recent advances in technology, mathematical tools and analytical solutions.

### 3.1. Empirical data and long-term monitoring programs: involving stakeholders and setting up fixed long-term monitoring stations across large spatial scales

In order to better understand zoonotic disease dynamics in wild animals we need reliable data sources to model spatial distribution and abundance of the host species involved in transmission, as good quality data is the cornerstone to building informed strategies that involve wildlife interventions. In reference to the badger-cattle bTB episystem in western Europe, both badgers (with several examples among the literature: [58, 59]), and cattle [60] have been extensively monitored. However, in some populations it is possible that deer and wild boar may also play a role in the local spread and maintenance of infection [61]. In Britain and Ireland, in the past decades, the significance of deer as a wildlife host impacting bTB epidemiology has been uncertain [62]. However, recent research is starting to uncover the role deer may play at local scales where conditions favour the transmission between badgers-deer-cattle [63]. There could be opportunities to gather data in collaboration with hunters (as has been used in France [64] and Spain [65], for example) to have access to an incredibly high number of deer samples, within and across countries (large spatial scales) and across years (long-term temporal scales). Involving stakeholders like hunters may provide the unique opportunity to use field tests for bTB [66, 67] to be submitted via smartphone applications, see [68], and to monitor the dynamic of the disease across multiple spatio-temporal scales and in relation to bTB occurrence in the two other hosts in the system (badgers and cattle). The ability to intercept the collaboration of stakeholders widely spread across large spatial scales (e.g., hunters, farmers, foresters) may help to establish systematic, relatively inexpensive, and long-term monitoring programmes able to provide data for Bayesian species and disease distribution models (described in section 3.3), allowing managers to access up to date risk scenarios. This approach can also highlight hotspots of disease outbreak that would drive very focused longitudinal studies involving satellite telemetry on multiple species simultaneously to better disentangle species overlaps and contact rates [69, 70]. The role of stakeholders/citizen scientists in this bTB example would be confirming infection, which is almost never economical, although there is the hope that cheaper field tests will be released in the next decade – or at least, we hope there will be investment in that direction. For the time being, a well distributed number of samples could be collected from hunters to cover large areas systematically and limit the costs required for testing.

When it comes to establishing long-term monitoring programs, fixed long-term sampling stations across large-spatial scales can capture wildlife population spatio-temporal dynamics. This can, on one hand, provide data on occurrence, relative density, and spatio-temporal overlaps of the host species and, on the other hand, gather key empirical data required to parameterise mathematical simulations. Camera traps are a popular and effective tool for estimating state variables of wildlife populations [71]. For ungulates, they have successfully been used to understand temporal behaviour (e.g., diel activity patterns, [72]), spatial behaviour (e.g., occupancy, [73]), and abundance (e.g., density, [74]). Camera traps have been used for quantifying temporal and spatial overlap of wild ungulates with domestic animals in open systems [75, 76] with varying results [77]. Kukielka *et al*. demonstrated their use in identifying hotspots of indirect wildlife–livestock overlap for the prevention of TB crossover [75]. For wildlife, especially ungulates, camera traps offer powerful monitoring solutions not only to measure abundance and spatial overlap, but also to understand behavioural dynamics that may align closely with disease risk. An example is the use of camera traps to individually recognise animals, which has been shown to be possible in a recent study by Hinojo *et al*. [78]. They demonstrated how roe deer (*Capreolus capreolus*) antler shapes could be used to identify distinct individuals. This data could be used to obtain better estimates of abundance as well as to build wildlife social networks (which will be discussed in more detail in section 3.3) to collect data on contact rates between and within species. The parameters from these analyses would be useful as an input for mathematical simulations to help better understand disease transmission dynamics in wildlife populations.

### 3.2. Recent advances with technology can help to gather data for mathematical simulations: interindividual variability within animal populations matters and should be taken into account

Detailed questions concerning the animal behaviour and movement patterns promoting disease transmission can be posed and answered using animal-attached sensors, i.e., biologging [79, 80]. GPS units are the most widely used of these sensors, providing data on animal locations and space use. Proximity sensors can detect when two or more sensor-equipped animals interact and can be used to detect direct encounters which may result in disease transmission. Collars with both GPS units and proximity sensors have been used concurrently on badgers and cattle, finding that while badgers show a habitat preference for cattle pastures, there were no direct contacts between the two species, suggesting that environmental transmission may play an important role in the case of bTB [81]. As such, proximity sensors allow insights which are not obtainable through investigating shared space use alone. Proximity sensors can also provide information on whether and how the duration of exposure to an infected individual affects transmission rate, where disease state of individuals is known [82]. Other biologging sensors, including accelerometers, magnetometers and gyroscopes, are used to classify distinct behaviours from logged datasets [80]. Behaviour classification allows activity budgets to be built, so that behaviours which increase the likelihood of acquiring or transmitting pathogens can be detected and mapped in the landscape. Accelerometers have also been used to compare micro-movements in diseased and healthy animals, with diseased animals exhibiting differences in posture, gait dynamism (e.g., the “bounce” in subsequent walking steps) and energy levels [83]. Monitoring such micro-movements in cattle could act as a warning sign to test herds for bTB when signs of illness are detected, by adapting existing systems in place to monitor lameness through accelerometry [84]. Further, these effects of disease on the internal state of animals yield important insights into how disease status impacts animal movement patterns and therefore disease spread.

Biologging and satellite telemetry monitoring can, on one hand, provide answers aimed at disentangling the transmission dynamics within multi-host disease systems [81, 85, 86] and, on the other hand, provide highly valuable empirical data that are strongly needed by parameter hungry mathematical simulations [82]. When tracking animals, however, special care should be spent to understand which animals we are monitoring, and whether we are following a special subset of individuals that are easier to trap. This applies also to where we study animals which will provide empirical data for mathematical simulations, because the behaviour and movement ecology may vary significantly depending on the level of human disturbance.

The concept of One Health has indeed highlighted the role of human activities in the spread of zoonotic diseases [87]. For example, urbanisation, improper waste disposal, and the intentional feeding of wildlife have been shown to result in wildlife movement into human-dominated areas [88], which may facilitate disease transfer to humans and other animal communities [89]. However, evidence has shown that only a select proportion of individuals within wildlife populations will engage in interactions with humans [90] or utilise these human-dominated areas [88, 91]. Individual variation in movement patterns [92], sociability [93] and immunological defence [94] among others, impacts disease transmission and spread [95]. There is supportive evidence that certain behavioural types have higher infection rates than others (e.g. [96, 97]), although the causal direction may be difficult to determine since infections also alter host behaviour [96, 98]. Regardless, to gain a more complete understanding of disease spread, future studies should incorporate this individual variation. These studies often utilise direct behavioural observations, since these are an invaluable data source that can be used to determine which individuals in a known population are more likely to engage in close-contact interactions with humans [90] or access human areas (e.g., farmland) [99]. This can provide us with information on which individuals in a population may be at “higher risk” of transferring disease to human or other animal populations.

### 3.3. Modelling and mathematical simulations: social network analysis, Bayesian species distribution models, and Agent Based Models

Social network analysis is a powerful tool for understanding the causes and consequences of disease transmission within the animal community [100, 101]. In the past decade social network analysis has mainly focused on understanding contact and transport networks of cattle and livestock movements, as well as wildlife movements [102-104]. Nonetheless, the area of network analysis can be expanded to better understand the dynamics of disease transmission between wildlife populations and livestock [101]. Unlike cattle monitoring the movements and interactions of wildlife can be challenging, where typically, a small proportion of individuals are monitored using recent advances with biologging and satellite telemetry, as discussed earlier. Recent advances in statistical analysis of social networks have paved the way to obtain better inferences from limited data [105-107]. Once the network metrics affecting disease transmission dynamics are identified (e.g., transitivity, betweenness centrality) [105, 106], those can be tested via pre-network permutations of available observations to ensure that the available data sufficiently captures non-random interactions among the animals. This can help estimate the scale of disease transmission in wild habitats. Global metrics of a social network help estimate the changes in the overall structure. For example, the network metric transitivity represents the tendency of a population to cluster together and is considered to be negatively correlated with disease transmission rates [104]. However, recent research [106] shows that it is noisily estimated at low sample sizes. Therefore, stable metrics with respect to low sample sizes should be identified before making inferences. Local network metrics such as betweenness centrality represent the tendency of an individual to serve as a bridge between one part of the community and another, helping the selection of individuals to be vaccinated/removed from the population. Research on data collected for wild ungulates [106] shows that the betweenness centrality values of smaller samples remain well correlated with those in larger samples. Similar correlation analysis can be done for other network metrics, mainly in cases of limited data availability for disease transmission. Whenever limited animals from a population are monitored, confidence intervals around the network metrics should also be obtained to make informed decisions using statistical evidence. Using the methodologies discussed by Silk *et al*. and Kaur *et al*. [104, 106] we now have the possibility of analysing all telemetry data collected thus far on species involved in bTB transmission (e.g., badgers, wild boar; but also applicable to species from other disease systems) to test hypotheses on disease transmission dynamics. In addition, it will help in collecting future data aimed at answering very precise questions such as understanding the role of deer species in bTB transmission by collecting simultaneous telemetry data of badgers and deer species in Ireland.

In reducing zoonotic risk, knowing the distribution and abundance of wildlife vectors is also essential [28, 108]. To that aim, Species Distribution Models (SDMs) can be used to produce models of the distribution and abundance of species based on occurrence data [109]. In recent years spatial modelling has undergone a conceptual and technical revolution. New modelling techniques within Bayesian [110] and Machine Learning frameworks [111] allow us to develop spatially explicit models of animal abundance and distributions with unprecedented accuracy, and the improvement of computational power allows computers to rise to the challenge and cope with the high computational demands of these models. The flexibility of the new techniques allows us to use different types of data (e.g., individual tracking data, survey data, and even citizen science data) and combine them in what is called Integrated Species Distribution Models (ISDMs), while still taking into account the different observational processes of each type of data, to produce accurate models even in data scarce systems [112]. In addition, these new techniques also allow for the calculation of uncertainty in a spatially explicit manner, which will help us evaluate the quality of the models and better interpret the results. Bayesian ISDMs using INLA (i.e., Integrated Nested Laplace Approximation) [113] were used to model the distribution of red, sika and fallow deer in Ireland, which are vectors of bTB [68]. The models produced, for the first time, relative abundance and distribution maps for each species, which will be an essential tool for deer population management and thus towards bTB eradication. They are already being used to determine high sika-density areas for a pilot study on the effect of deer on biodiversity, which will provide further management tools for the overabundant deer populations in Ireland. In addition, hierarchical Bayesian models are also the basis of a new project aimed at modelling European badger sett distribution, badger density, and their body condition. These three models will be linked to bTB infection in badgers and outbreaks in cattle, in an attempt for the first time to link badger spatial ecology to bTB management and eradication in Ireland (V. Morera-Pujol 2023, personal communication).

Agent-based simulations are a powerful tool for understanding the transient effects of wildlife disease in human-dominated landscapes, both as a complementary tool to traditional methods and as a surrogate for scenarios where data is limited or not available [16]. These models serve as a computational laboratory that allow researchers to plug-in available real-world data and parameterise both agents (for instance, a badger) and the environment (for instance, a mosaic of natural habitats and farms) to empirically test if animal behaviour in response to landscape change or management interventions modulates disease risk dynamics over time and space [114]. Recent technological advancements have bolstered agent-based simulations allowing for high-resolution spatio-temporal models that incorporate geographic information systems (GIS) data to create hyper realistic environments, and machine learning algorithms to introduce cognition and applied decision making for agents. Furthermore, process-driven agent-based models (e.g., disease transmission) can be integrated into larger mechanistic agent-based models (e.g., ecosystem scale epi-dynamics) for increasingly higher-resolution models that reduce uncertainty and overly-theoretical parameterisation of model entities [16]. The development of hyper-realistic agent-based simulations, parameterised with high-resolution data, for the management of bovine tuberculosis in multi-host systems can contribute to answering important policy questions and how best to select management directions. In practice, this allows for the totality of data collected in complex multi-host systems to be incorporated into a single environment where they may be measured against one another in the simulation to deduce the possible effects of each predictor. Take for example the European badger as the primary wildlife host in Ireland as a case study. Badgers are prevalent in the agroecological mosaic of natural habitats and farms in Ireland. Agent-based simulations can utilise data from badger tracking studies [53, 115, 116], habitat suitability [117], culling and vaccination programmes [118] and disturbance regimes [54, 119] to simulate badger movement and behaviour realistically. GIS data for farm location and characteristics [120], as well as ecological and environmental data streams, can then place the badger agent into a highly realistic environment to examine how these factors affect badger movement, behaviour, and other parameters, for instance, contact rates with domestic animals. Interactions between agents and the environment can be modulated by sub-models to further increase the strength of the model, for instance, weather sub-models (e.g., rainfall) may influence agricultural practice and thus contact rates, alternatively, disease transmission could also be sub-modelled so that contact rates may/may not result in infection [16]. Finally, management decisions can be trialled within the simulation to see how likely decisions change the status of disease within the study system, allowing for “What if?” scenarios to play out without risk to animal or human welfare or livelihoods.

Our review is a starting point to better understand the current gaps in the zoonotic disease literature. We are aware that this review has some limitations such as the use of only one search engine, as well as focusing on one zoonotic disease system and date range. However, we hope that it will incite new and exciting possibilities for the future which could expand on this current work, by for example looking at multiple disease systems as well as other date ranges to try and uncover more information on possible way forwards to combat zoonotic disease transmission.

## Conclusion

Our exploration of the recent literature on multi-host bTB episystems, as an example of zoonotic One Health challenges, has revealed a significant body of work utilising a diversity of methodologies at different spatio-temporal scales and subjects (individual vs. group) levels. There was a significant bias in the literature towards one particular episystem, the badger-cattle system that predominates in north-western Europe, reflecting large financial burdens (for both governments as well as the agricultural sector) and research funding investments. Alternatively, there were comparatively less publications from the global south, especially in complex muti-host episystems in southern Africa. In such episystems, the cost-effective and efficient collection, collation, and use of data are essential to drive greatest added value to inform on policy options. A theme from the present work is the essential need to use new and emerging technologies to derive new insights from data that may have been collected for other purposes.

Given the results from our scoping review, we reflect on several areas where progress could be made, including the need for high-quality data on wildlife hosts, even in episystems where significant research investments have already been made. Such careful collection and utilisation of empirical data could then feed the development of social network analyses, Bayesian distribution models and eventually Agent Based Models. Mathematical simulations such as ABMs trained on synthetic data and parameterised by real empirical data can answer questions that would otherwise be too costly, unethical, or both. Such models can also be used to explore different scenarios in an increasingly human-dominated world, under different levels of land-use and climate change, or with the appearance of invasive species in already complex multi-host epidemiological systems. In addition, it can help build cross-disciplinary bridges with other areas, deriving significant insights into interspecific transmission like phylodynamic modelling.

Taking Ireland as an example, with the intention not to stay local but rather inspire researchers from across the globe; Ireland invests considerably in surveying, culling, and vaccinating badgers [121]. However, the question remains - which applies to other countries and zoonotic episystems - should we be doing more, or being smarter with the data we already have? We suggest the latter. Yes, we need to be smarter, arranging *ad hoc* data collections using the latest technological tools to estimate unknown parameters. We need to focus our efforts on mathematical modelling (ABMs, INLA-Bayesian) to optimise our information gain from the large, high-quality datasets collected over the last few decades (and for sparser datasets, taking advantage of recently developed statistical tools for enhanced inferences, see [54, 68]). We have (almost) all the data required to parameterise simulations with significant utility: this should be one main focus in future research. We believe that, ideally, the feedback of simulation and mathematical tools to inform data collection, and the “virtuous cycle” of feeding this new data to improve the next generation of model is a priority for decision making tools for policy makers and programme managers.

## Declarations

### Ethics approval and consent to participate

Not applicable

### Consent for publication

Not applicable

### Availability of data and materials

The datasets generated and/or analysed during the current study are available in the ‘*Curbing zoonotic disease spread in multi-host-species systems will require integrating novel data streams and analytical approaches: evidence from a scoping review of bovine tuberculosis*’ repository, https://doi.org/10.5281/zenodo.7886368

### Competing interests

The authors declare that they have no competing interests.

## Funding

This publication has emanated from research conducted with the financial support of Science Foundation Ireland under Grant number 18/CRT/6049. For the purpose of Open Access, the author has applied a CC BY public copyright licence to any Author Accepted Manuscript version arising from this submission.

In addition, HME is funded by an Irish Research Council Government of Ireland postgraduate scholarship.

### Authors’ contributions

KC - conceived and designed the review, acquired, analysed and interpreted the data, wrote the first draft of the manuscript.

HME - edited and revised the ms and contributed to the interpretation of the data.

AWB - edited and revised the ms and contributed to the interpretation of the data, drafted the policy and research implications.

BA, LLG, PK, VMP, KJM, AFS, MST - edited specific sections of the discussion and revised the whole ms. SC - conceived and designed the review, edited the ms and supervised the process.

All Authors approved the final version of this review.

## Acknowledgements

We would like to acknowledge Sylvia Power for helping in reviewing the final version of the manuscript.

## Notes

https://doi.org/10.5281/zenodo.7886368

### Competing Interest Statement

The authors have declared no competing interest.

### Summary of Updates

Abstract format updated

