## Additional file 1 for "Curbing zoonotic disease spread in multi-host-species systems will require integrating novel data streams and analytical approaches: evidence from a scoping review of bovine tuberculosis" (from another version of this preprint: version 1, 2023-05-10)

**Web of Science Search Terms:**

TS=((cattle OR livestock OR badger OR deer OR elk OR boar OR "feral pig" OR "wild pig" OR possum OR buffalo)AND ("bovine tuberculosis" OR "bovine TB" OR "mycobacterium bovis" OR "M. bovis") AND (r isk* OR management OR control* OR presence OR prevalence OR occurrence OR spread* OR transmis sion)) Indexes=SCI-EXPANDED, SSCI, A&HCI, CPCI-S, CPCI-SSH, BKCI-S, BKCI-SSH, ESCI, CCR-EXPANDED, IC Timespan=All years

**The search above was completed on the following dates:**

- 11/3/2021
- 20/7/2022

**Below can be found the list of articles selected as relevant [1-539]. Articles included in the results were articles published between January 2017 to July 2022 (shown in bold).**


1. **Abbate JM, Arfuso F, Iaria C, Arestia G, Lanteri G (2020) Prevalence of Bovine Tuberculosis in Slaughtered Cattle in Sicily, Southern Italy. Animals 10(9):11.**
2. **Abdelaal HFM, Spalink D, Amer A, Steinberg H, Hashish EA, Nasr EA, Talaat AM (2019) Genomic Polymorphism Associated with the Emergence of Virulent Isolates of Mycobacterium bovis in the Nile Delta. Scientific Reports 9:15.**
3. Abdou M, Frankena K, O'Keeffe J, Byrne AW (2016) Effect of culling and vaccination on bovine tuberculosis infection in a European badger (Meles meles) population by spatial simulation modelling. Preventive Veterinary Medicine 125:19-30.
4. Abernethy DA, Denny GO, Menzies FD, McGuckian P, Honhold N, Roberts AR (2006) The Northern Ireland programme for the control and eradication of Mycobacterium bovis. Veterinary Microbiology 112(2-4):231-237.
5. Abernethy DA, Upton P, Higgins IM, McGrath G, Goodchild AV, Rolfe SJ, Broughan JM, Downs SH, Clifton-Hadley R, Menzies FD et al (2013) Bovine tuberculosis trends in the UK and the Republic of Ireland, 1995-2010. Veterinary Record 172(12):14.
6. **Abrantes AC, Serejo J, Vieira-Pinto M (2021) The Association between Palmer Drought Severity Index Data and Tuberculosis-like Lesions Occurrence in Mediterranean Hunted Wild Boars. Animals 11(7):8.**
7. **Acevedo P, Prieto M, Quiros P, Merediz I, de Juan L, Infantes-Lorenzo JA, Triguero-Ocana R, Balseiro A (2019) Tuberculosis Epidemiology and Badger (Meles meles) Spatial Ecology in a Hot-Spot Area in Atlantic Spain. Pathogens 8(4):18.**
8. Acevedo P, Romero B, Vicente J, Caracappa S, Galluzzo P, Marineo S, Vicari D, Torina A, Casal C, de la Fuente J et al (2013) Tuberculosis Epidemiology in Islands: Insularity, Hosts and Trade. Plos One 8(7):9.
9. Adkin A, Brouwer A, Simons RRL, Smith RP, Arnold ME, Broughan J, Kosmider R, Downs SH (2016) Development of risk-based trading farm scoring system to assist with the control of bovine tuberculosis in cattle in England and Wales. Preventive Veterinary Medicine 123:32-38.
10. Agusto FB, Lenhart S, Gumel AB, Odoi A (2011) Mathematical analysis of a model for the transmission dynamics of bovine tuberculosis. Mathematical Methods in the Applied Sciences 34(15):1873-1887.
11. **Alberti TS, Bruhn FRP, Zamboni R, Venancio FR, Scheid HV, Raffi MB, Schild AL, Sallis ESV (2020) Epidemiological analysis of bovine tuberculosis in the southern region of Rio Grande do Sul from 2000 to 2015. Pesquisa Veterinaria Brasileira 40(2):77-81.**
12. Alberto JR, Aranha JM, Serejo JP, Amado A, Vieira-Pinto M (2011) The utility of GIS in studying the distribution of Bovine Tuberculosis in wild boar (Sus scrofa) and red deer (Cervus elaphus) in Central Portugal.

progression of tuberculosis in badgers. Proceedings of the Royal Society B-Biological Sciences 278(1713):1913-1920.
98. Che'Amat A, Armenteros JA, Gonzalez-Barrio D, Lima JF, Diez-Delgado I, Barasona JA, Romero B, Lyashchenko KP, Ortiz JA, Gortazar C (2016) Is targeted removal a suitable means for tuberculosis control in wild boar? Preventive Veterinary Medicine 135:132-135.
99. Cheeseman CL, Jones GW, Gallagher J, Mallinson PJ (1981) The population-structure, density and prevalence of tuberculosis (mycobacterium-bovis) in badgers (meles-meles) from 4 areas in southwest england. Journal of Applied Ecology 18(3):795-804.
**100. Ciaravino G, Garcia-Saenz A, Cabras S, Allepuz A, Casal J, Garcia-Bocanegra I, De Koeijer A, Gubbins S, Saez JL, Cano-Terriza D et al (2018) Assessing the variability in transmission of bovine tuberculosis within Spanish cattle herds. Epidemics 23:110-120.**
**101. Ciaravino G, Laranjo-Gonzalez M, Casal J, Saez-Llorente JL, Allepuz A (2021) Most likely causes of infection and risk factors for tuberculosis in Spanish cattle herds. Veterinary Record 189(2):11.**
**102. Clarke A, Byrne AW, Maher J, Ryan E, Farrell F, McSweeney C, Barrett D (2022) Engaging With Farmers to Explore Correlates of Bovine Tuberculosis Risk in an Internationally Important Heritage Landscape: The Burren, in the West of Ireland. Frontiers in Veterinary Science 9:9.**
103. Cleaveland S, Shaw DJ, Mfinanga SG, Shirima G, Kazwala RR, Eblate E, Sharp M (2007) Mycobacterium bovis in rural Tanzania: Risk factors for infection in human and cattle populations. Tuberculosis 87(1):30-43.
104. Clegg TA, Blake M, Healy R, Good M, Higgins IM, More SJ (2013) The impact of animal introductions during herd restrictions on future herd-level bovine tuberculosis risk. Preventive Veterinary Medicine 109(3-4):246-257.
**105. Clegg TA, Good M, Hayes M, Duignan A, McGrath G, More SJ (2018) Trends and Predictors of Large Tuberculosis Episodes in Cattle Herds in Ireland. Frontiers in Veterinary Science 5:12.**
106. Clegg TA, Good M, More SJ (2016) Risk factors for cattle presenting with a confirmed bTB lesion at slaughter, from herds with no evidence of within-herd transmission. Preventive Veterinary Medicine 126:111-120.
107. Cliftonhadley RS, Wilesmith JW, Richards MS, Upton P, Johnston S (1995) The occurrence of mycobacterium-bovis infection in cattle in and around an area subject to extensive badger (meles-meles) control. Epidemiology and Infection 114(1):179-193.
108. Coleman JD, Coleman MC, Warburton B (2006) Trends in the incidence of tuberculosis in possums and livestock, associated with differing control intensities applied to possum populations. New Zealand Veterinary Journal 54(2):52-60.
109. Coleman JD, Cooke MM, Jackson R, Webster R (1999) Temporal patterns in bovine tuberculosis in a brushtail possum population contiguous with infected cattle in the Ahaura Valley, Westland. New Zealand Veterinary Journal 47(4):119-124.
110. Coleman JD, Jackson R, Cooke MM, Grueber L (1994) Prevalence and spatial-distribution of bovine tuberculosis in brushtail possums on a forest-scrub margin. New Zealand Veterinary Journal 42(4):128-132.
111. Conlan AJK, McKinley TJ, Karolemeas K, Pollock EB, Goodchild AV, Mitchell AP, Birch CPD, Clifton-Hadley RS, Wood JLN (2012) Estimating the Hidden Burden of Bovine Tuberculosis in Great Britain. Plos Computational Biology 8(10):14.
112. Cook AJC, Tuchili LM, Buve A, Foster SD, GodfreyFaussett P, Pandey GS, McAdam K (1996) Human and bovine tuberculosis in the Monze District of Zambia - A cross-sectional study. British Veterinary Journal 152(1):37-46.
113. Corn JL, Nettles VF (2001) Health protocol for translocation of free-ranging elk. Journal of Wildlife Diseases 37(3):413-426.

147. Donnelly CA, Woodroffe R, Cox DR, Bourne J, Gettinby G, Le Fevre AM, McInerney JP, Morrison WI (2003) Impact of localized badger culling on tuberculosis incidence in British cattle. Nature 426(6968):834-837.
**148. Downs SH, Ashfield S, Arnold M, Roberts T, Prosser A, Robertson A, Frost S, Harris K, Avigad R, Smith GC (2022) Detection of a local Mycobacterium bovis reservoir using cattle surveillance data. Transboundary and Emerging Diseases:15.**
**149. Doyle LP, Courcier EA, Gordon AW, O'Hagan MJH, Stegeman JA, Menzies FD (2017) Bovine tuberculosis in Northern Ireland: quantification of the population disease-level effect from cattle leaving herds detected as a source of infection. Epidemiology and Infection 145(16):3505-3515.**
150. Drewe JA, O'Connor HM, Weber N, McDonald RA, Delahay RJ (2013) Patterns of direct and indirect contact between cattle and badgers naturally infected with tuberculosis. Epidemiology and Infection 141(7):1467-1475.
**151. Duault H, Michelet L, Boschiroli ML, Durand B, Canini L (2022) A Bayesian evolutionary model towards understanding wildlife contribution to F4-family Mycobacterium bovis transmission in the South-West of France. Veterinary Research 53(1):12.**
152. Echeverria G, Ron L, Leon AM, Espinosa W, Benitez-Ortiz W, Proano-Perez F (2014) Prevalence of bovine tuberculosis in slaughtered cattle identified by nested-PCR in abattoirs from two dairy areas of Ecuador. Tropical Animal Health and Production 46(6):1015-1022.
153. Egbe NF, Muwonge A, Ndip L, Kelly RF, Sander M, Tanya V, Ngwa VN, Handel IG, Novak A, Ngandalo R et al (2016) Abattoir-based estimates of mycobacterial infections in Cameroon. Scientific Reports 6:14.
154. Elias K, Hussein D, Asseged B, Wondwossen T, Gebeyehu M (2008) Status of bovine tuberculosis in Addis Ababa dairy farms. Revue Scientifique Et Technique-Office International Des Epizooties 27(3):915-923.
155. Enriquez-Cruz C, Cruz-Hernandez NI, Zertuche-Rodriguez JL, Uriegas-Garcia JL, Toscano-Ruiz JE, Flores-Gutierrez GH (2010) Epidemiology of bovine tuberculosis in Mexico, bordering the United States, at establishment of controlling strategies. Arquivo Brasileiro De Medicina Veterinaria E Zootecnia 62(5):1029-1035.
**156. Escarcega DAV, Razo CAP, Ruiz SG, Gallegos SLS, Suazo FM, Alarcon GJC (2020) Analysis of bovine tuberculosis transmission in Jalisco, Mexico through whole-genome sequencing. Journal of Veterinary Research 64(1):51-61.**
**157. Esmaeilzadeh N, Bahonar A, Foroushani AR, Nasehi M, Amiri K, Hadjzadeh MAR (2022) Temporal trends and prediction of bovine tuberculosis: a time series analysis in the North-East of Iran. Iranian Journal of Veterinary Research 23(1):12-17.**
158. Fenichel EP, Horan RD (2007) Gender-Based harvesting in wildlife disease management. American Journal of Agricultural Economics 89(4):904-920.
159. Fenichel EP, Horan RD (2007) Jointly-determined ecological thresholds and economic trade-offs in wildlife disease management. Natural Resource Modeling 20(4):511-547.
160. Fenwick NID (2012) Modelled impacts of badger culling on cattle TB in a real area with geographic boundaries. Veterinary Record 170(7):177-+.
161. Fetene T, Kebede N (2009) Bovine tuberculosis of cattle in three districts of northwestern Ethiopia. Tropical Animal Health and Production 41(2):273-277.
**162. Fielding HR, McKinley TJ, Delahay RJ, Silk MJ, McDonald RA (2020) Effects of trading networks on the risk of bovine tuberculosis incidents on cattle farms in Great Britain. Royal Society Open Science 7(4):11.**
**163. Fielding HR, McKinley TJ, Silk MJ, Delahay RJ, McDonald RA (2019) Contact chains of cattle farms in Great Britain. Royal Society Open Science 6(2):16.**
**164. Fielding HR, Silk MJ, McKinley TJ, Delahay RJ, Wilson-Aggarwal JK, Gauvin L, Ozella L, Cattuto C, McDonald RA (2021) Spatial and temporal variation in proximity networks of commercial dairy cattle in Great Britain. Preventive Veterinary Medicine 194:11.**

**270. Mareledwane VE, Adesiyun AA, Thompson PN, Hlokwe TM (2021) Application of the gamma-interferon assay to determine the prevalence of bovine tuberculosis in slaughter livestock at abattoirs in Gauteng, South Africa. Veterinary Medicine and Science:8.**

283. McInerney J, Small KJ, Caley P (1995) Prevalence of Mycobacterium bovis infection in feral pigs in the Northern Territory. Australian Veterinary Journal 72(12):448-451.
284. McKenzie JS, Morris RS, Pfeiffer DU, Dymond JR (2002) Application of remote sensing to enhance the control of wildlife-associated Mycobacterium bovis infection. Photogrammetric Engineering and Remote Sensing 68(2):153-159.
**285. McKinley TJ, Lipschutz-Powell D, Mitchell AP, Wood JLN, Conlan AJK (2018) Risk factors and variations in detection of new bovine tuberculosis breakdowns via slaughterhouse surveillance in Great Britain. Plos One 13(6):14.**
**286. Mekonnen GA, Ameni G, Wood JLN, Berg S, Conlan AJK, Aseffa A, Mihret A, Tessema B, Belachew B, Fekadu EW et al (2019) Network analysis of dairy cattle movement and associations with bovine tuberculosis spread and control in emerging dairy belts of Ethiopia. Bmc Veterinary Research 15:14.**
**287. Mekonnen GA, Conlan AJK, Berg S, Ayele BT, Alemu A, Guta S, Lakew M, Tadesse B, Gebre S, Wood JLN et al (2019) Prevalence of bovine tuberculosis and its associated risk factors in the emerging dairy belts of regional cities in Ethiopia. Preventive Veterinary Medicine 168:81-89.**
**288. Mekonnen GA, Conlan AJK, Berg S, Ayele BT, Mihret A, Olani A, Asgedom H, Aseffa A, Wood JLN, Ameni G (2021) Dynamics and risk of transmission of bovine tuberculosis in the emerging dairy regions of Ethiopia. Epidemiology and Infection 149:9.**
289. Mentaberre G, Romero B, de Juan L, Navarro-Gonzalez N, Velarde R, Mateos A, Marco I, Olive-Boix X, Dominguez L, Lavin S et al (2014) Long-Term Assessment of Wild Boar Harvesting and Cattle Removal for Bovine Tuberculosis Control in Free Ranging Populations. Plos One 9(2):12.
290. Menzies FD, Abernethy DA, Stringer LA, Honhold N, Gordon AW (2012) A matched cohort study investigating the risk of Mycobacterium bovis infection in the progeny of infected cows. Veterinary Journal 194(3):299-302.
291. Menzies FD, Abernethy DA, Stringer LA, Jordan C (2011) A comparison of badger activity in two areas of high and low bovine tuberculosis incidence of Northern Ireland. Veterinary Microbiology 151(1-2):112-119.
**292. Menzies FD, McCormick CM, O'Hagan MJH, Collins SF, McEwan J, McGeown CF, McHugh GE, Hart CD, Stringer LA, Molloy C et al (2021) Test and vaccinate or remove: Methodology and preliminary results from a badger intervention research project. Veterinary Record 189(5):12.**
**293. Meunier NV, Sebulime P, White RG, Kock R (2017) Wildlife-livestock interactions and risk areas for cross-species spread of bovine tuberculosis. Onderstepoort Journal of Veterinary Research 84(1):10.**
294. Milian-Suazo F, Salman MD, Ramirez C, Payeur JB, Rhyan JC, Santillan M (2000) Identification of tuberculosis in cattle slaughtered in Mexico. American Journal of Veterinary Research 61(1):86-89.
295. Mill AC, Rushton SP, Shirley MDF, Murray AWA, Smith GC, Delahay RJ, McDonald RA (2012) Farm-scale risk factors for bovine tuberculosis incidence in cattle herds during the Randomized Badger Culling Trial. Epidemiology and Infection 140(2):219-230.
296. Miller R, Kaneene JB, Fitzgerald SD, Schmitt SM (2003) Evaluation of the influence of supplemental feeding of white-tailed deer (Odocoileus virginianus) on the prevalence of bovine tuberculosis in the Michigan wild deer population. Journal of Wildlife Diseases 39(1):84-95.
297. Miller R, Kaneene JB, Schmitt SM, Lusch DP, Fitzgerald SD (2004) Geographic distribution and spatial analysis of Mycobacterium bovis in Michigan white-tailed deer.
298. Miller R, Kaneene JB, Schmitt SM, Lusch DP, Fitzgerald SD (2007) Spatial analysis of Mycobacterium bovis infection in white-tailed deer (Odocoileus virginianus) in Michigan, USA. Preventive Veterinary Medicine 82(1-2):111-122.

315. Munyeme M, Muma JB, Samui KL, Skjerve E, Nambota AM, Phiri IGK, Rigouts L, Tryland M (2009) Prevalence of bovine tuberculosis and animal level risk factors for indigenous cattle

under different grazing strategies in the livestock/wildlife interface areas of Zambia. Tropical Animal Health and Production 41(3):345-352.

316. Munyeme M, Muma JB, Skjerve E, Nambota AM, Phiri IGK, Samui KL, Dorny P, Tryland M (2008) Risk factors associated with bovine tuberculosis in traditional cattle of the livestock/wildlife interface areas in the Kafue basin of Zambia. Preventive Veterinary Medicine 85(3-4):317-328.

**317. Murphy KJ, Morera-Pujol V, Ryan E, Byrne AW, Breslin P, Ciuti S (2022) Habitat availability alters the relative risk of a bovine tuberculosis breakdown in the aftermath of a commercial forest clearfell disturbance. Journal of Applied Ecology:13.**

347. Oloya J, Muma JB, Opuda-Asibo J, Djonne B, Kazwala R, Skjerve E (2007) Risk factors for herd-level bovine-tuberculosis seropositivity in transhumant cattle in Uganda. Preventive Veterinary Medicine 80(4):318-329.
348. Oloya J, Opuda-Asibo J, Djonne B, Muma JB, Matope G, Kazwala R, Skjerve E (2006) Responses to tuberculin among Zebu cattle in the transhumance regions of Karamoja and Nakasongola district of Uganda. Tropical Animal Health and Production 38(4):275-283.
349. Omer MK, Skjerve E, Woldehiwet Z, Holstad G (2001) A cross-sectional study of bovine tuberculosis in dairy farms in Asmara, Eritrea. Tropical Animal Health and Production 33(4):295-303.
**350. Orrico M, van Schaik G, Koets A, van den Broek J, Montizaan M, La Haye M, Rijks JM (2022) The effectiveness of bovine tuberculosis surveillance in Dutch badgers. Transboundary and Emerging Diseases:13.**
**351. Orton RJ, Deason M, Bessell PR, Green DM, Kao RR, Salvador LCM (2018) Identifying genotype specific elevated-risk areas and associated herd risk factors for bovine tuberculosis spread in British cattle. Epidemics 24:34-42.**
352. Palisson A, Courcoul A, Durand B (2016) Role of Cattle Movements in Bovine Tuberculosis Spread in France between 2005 and 2014. Plos One 11(3):19.
353. Palmer MV, Waters WR, Whipple DL (2002) Milk containing Mycobacterium bovis as a source of infection for white-tailed deer fawns (Odocoileus virginianus). Tuberculosis 82(4-5):161-165.
354. Palmer MV, Waters WR, Whipple DL (2004) Investigation of the transmission of Mycobacterium bovis from deer to cattle through indirect contact. American Journal of Veterinary Research 65(11):1483-1489.
355. Palmer MV, Waters WR, Whipple DL (2004) Shared feed as a means of deer-to-deer transmission of Mycobacterium bovis. Journal of Wildlife Diseases 40(1):87-91.
356. Palmer MV, Whipple DL, Waters WR (2001) Experimental deer-to-deer transmission of Mycobacterium bovis. American Journal of Veterinary Research 62(5):692-696.
357. Parra A, Garcia A, Inglis NF, Tato A, Alonso JM, de Mendoza MH, de Mendoza JH, Larrasa J (2006) An epidemiological evaluation of Mycobacterium bovis infections in wild game animals of the Spanish Mediterranean ecosystem. Research in Veterinary Science 80(2):140-146.
**358. Pascual-Linaza AV, Gordon AW, Stringer LA, Menzies FD (2017) Efficiency of slaughterhouse surveillance for the detection of bovine tuberculosis in cattle in Northern Ireland. Epidemiology and Infection 145(5):995-1005.**
359. Paterson BM, Morris RS (1995) Interactions between beef cattle and simulated tuberculous possums on pasture. New Zealand Veterinary Journal 43(7):289-293.
360. Paterson BM, Morris RS, Weston J, Cowan PE (1995) Foraging and denning patterns of brushtail possums, and their possible relationship to contact with cattle and the transmission of bovine tuberculosis. New Zealand Veterinary Journal 43(7):281-288.
361. Pavlik I, Ayele WY, Parmova I, Melicharek I, Hanzlikova M, Kormendy B, Nagy G, Cvetnic Z, Ocepek M, Fejzic N et al (2002) Incidence of bovine tuberculosis in cattle in seven Central European countries during the years 1990-1999. Veterinarni Medicina 47(2-3):45-51.
362. Pavlik I, Trcka I, Parmova I, Svobodova J, Melicharek I, Nagy G, Cvetnic Z, Ocepek M, Pate M, Lipiec M (2005) Detection of bovine and human tuberculosis in cattle and other animals in six Central European countries during the years 2000-2004. Veterinarni Medicina 50(7):291-299.
363. Payne A, Boschiroli ML, Gueneau E, Moyen JL, Rambaud T, Dufour B, Gilot-Fromont E, Hars J (2013) Bovine tuberculosis in "Eurasian" badgers (Meles meles) in France. European Journal of Wildlife Research 59(3):331-339.
364. Payne A, Chappa S, Hars J, Dufour B, Gilot-Fromont E (2016) Wildlife visits to farm facilities assessed by camera traps in a bovine tuberculosis-infected area in France. European Journal of Wildlife Research 62(1):33-42.

**465. Tanner E, White A, Lurz PWW, Gortazar C, Diez-Delgado I, Boots M (2019) The Critical Role of Infectious Disease in Compensatory Population Growth in Response to Culling. American Naturalist 194(1):E1-E12.**
466. Tasmi, Aldila D, Soewono E, Nuraini N (2016) Mathematical Model for Transmission of Tuberculosis in Badger Population with Vaccination. In:Edited by Takeuchi Y, Apri M (eds) Symposium on Biomathematics, Volume 1723, edn.
467. Thompson AK, Samuel MD, Van Deelen TR (2008) Alternative feeding strategies and potential disease transmission in Wisconsin white-tailed deer. Journal of Wildlife Management 72(2):416-421.
468. Tigre W, Alemayehu G, Abetu T, Ameni G (2012) Preliminary study on the epidemiology of bovine tuberculosis in Jimma town and its surroundings, Southwestern Ethiopia. African Journal of Microbiology Research 6(11):2591-2597.
469. Tipu MY, Chaudhary ZI, Younus M, Rabbani M (2012) A Cross Sectional Study of Mycobacterium bovis in Dairy Cattle in and Around Lahore City, Pakistan. Pakistan Journal of Zoology 44(2):393-398.
470. Tolhurst BA, Delahay RJ, Walker NJ, Ward AI, Roper TJ (2009) Behaviour of badgers (Meles meles) in farm buildings: Opportunities for the transmission of Mycobacterium bovis to cattle? Applied Animal Behaviour Science 117(1-2):103-113.
471. Tolhurst BA, Ward AI, Delahay RJ, MacMaster AM, Roper TJ (2008) The behavioural responses of badgers (Meles meles) to exclusion from farm buildings using an electric fence. Applied Animal Behaviour Science 113(1-3):224-235.
472. Tomlinson AJ, Chambers MA, Carter SP, Wilson GJ, Smith GC, McDonald RA, Delahay RJ (2013) Heterogeneity in the risk of Mycobacterium bovis infection in European badger (Meles meles) cubs. Epidemiology and Infection 141(7):1458-1466.
473. Tompkins DM, Ramsey DSL, Cross ML, Aldwell FE, de Lisle GW, Buddle BM (2009) Oral vaccination reduces the incidence of tuberculosis in free-living brushtail possums. Proceedings of the Royal Society B-Biological Sciences 276(1669):2987-2995.
**474. Tosa MI, Schauber EM, Nielsen CK (2017) Localized removal affects white-tailed deer space use and contacts. Journal of Wildlife Management 81(1):26-37.**
**475. Tratalos JA, Madden JM, McGrath G, Graham DA, Collins AB, More SJ (2020) Spatial and network characteristics of Irish cattle movements. Preventive Veterinary Medicine 183:17.**
476. Trewby H, Wright D, Breadon EL, Lycett SJ, Mallon TR, McCormick C, Johnson P, Orton RJ, Allen AR, Galbraith J et al (2016) Use of bacterial whole-genome sequencing to investigate local persistence and spread in bovine tuberculosis. Epidemics 14:26-35.
**477. Triguero-Ocana R, Laguna E, Jimenez-Ruiz S, Fernandez-Lopez J, Garcia-Bocanegra I, Barasona JA, Risalde MA, Montoro V, Vicente J, Acevedo P (2021) The wildlife-livestock interface on extensive free-ranging pig farms in central Spain during the "montanera" period. Transboundary and Emerging Diseases 68(4):2066-2078.**
**478. Triguero-Ocana R, Martinez-Lopez B, Vicente J, Barasona JA, Martinez-Guijosa J, Acevedo P (2020) Dynamic Network of Interactions in the Wildlife-Livestock Interface in Mediterranean Spain: An Epidemiological Point of View. Pathogens 9(2):16.**
479. Tschopp R, Bekele S, Moti T, Young D, Aseffa A (2015) Brucellosis and bovine tuberculosis prevalence in livestock from pastoralist communities adjacent to Awash National Park, Ethiopia. Preventive Veterinary Medicine 120(2):187-194.
480. Tschopp R, Schelling E, Hattendorf J, Aseffa A, Zinsstag J (2009) Risk factors of bovine tuberculosis in cattle in rural livestock production systems of Ethiopia. Preventive Veterinary Medicine 89(3-4):205-211.
481. Tschopp R, Schelling E, Hattendorf J, Young D, Aseffa A, Zinsstag J (2010) Repeated cross-sectional skin testing for bovine tuberculosis in cattle kept in a traditional husbandry system in Ethiopia. Veterinary Record 167(7):250-256.
